# Multi-omic profiling of squamous cell lung cancer identifies metabolites and related genes associated with squamous cell carcinoma

**DOI:** 10.1101/2024.04.17.589879

**Authors:** Johan Staaf, Daniel Ehinger, Hans Brunnström, Mats Jönsson, Frida Rosengren, Marija Kotevska, Anna Karlsson, Mattias Aine, Christian Frezza, Maria Planck, Elsa Arbajian

**Affiliations:** Division of Oncology, Department of Clinical Sciences Lund, Lund University, Medicon Village, SE-22381 Lund, Sweden; Division of Translational Cancer Research, Department of Laboratory medicine, Lund University, Medicon Village, SE-22381 Lund, Sweden; Department of Genetics, Pathology, and Molecular Diagnostics, Skåne University Hospital, Lund, Sweden; Division of Pathology, Department of Clinical Sciences Lund, Lund University, Lund, Sweden; Division of Respiratory Medicine, Allergology, and Palliative Medicine, Department of Clinical Sciences Lund, Lund University, Skåne University Hospital, SE-221 85 Lund, Sweden; University of Cologne, Faculty of Medicine and University Hospital Cologne, Institute for Metabolomics in Ageing, Cluster of Excellence Cellular Stress Responses in Aging-associated Diseases (CECAD), Cologne, Germany; University of Cologne, Faculty of Mathematics and Natural Sciences, Institute of Genetics, Cluster of Excellence Cellular Stress Responses in Aging-associated Diseases (CECAD), Cologne, Germany

**Keywords:** Lung cancer, squamous cell carcinoma, metabolomics, SLC6A8, creatine

## Abstract

**Background:** Squamous cell lung carcinoma (SqCC) is the second most common histological subtype of lung cancer. Besides -tumor-initiating and promoting DNA, RNA, and epigenetic alterations, aberrant tumor cell metabolism has been identified as one of the hallmarks of carcinogenesis. The aim of the current study was to identify SqCC-specific metabolites and key gene regulators that could eventually be used as new anticancer targets.

**Methods:** Transcriptional (n=156), proteomics (n=118), and mass spectrometry-based metabolomic data (n=73) were gathered for a cohort of resected early-stage lung cancers representing all major histological subgroups. SqCC-specific differentially expressed genes were integrated with proteogenomic and metabolic data using genome scale metabolic models (GEMs). Findings were validated in cohorts of tumors, normal specimens, and cancer cell lines. In situ protein expression of SLC6A8 was investigated in 213 tumors.

**Results:** Differential gene expression analysis identified 280 SqCC-specific genes, of which 57 were connected to metabolites through GEMs. Metabolic profiling identified 7 SqCC-specific metabolites, of which increased creatine and decreased phosphocholine levels matched to SqCC-specific elevated expression of *SLC6A8* and decreased expression of *CHKA,* part of respective GEMs. Expression of both genes appeared tumor cell-associated, and in particular the elevated expression of *SLC6A8* identified SqCC also in stage IV disease.

**Conclusion:** Elevated creatine levels and the overexpression of its transporter protein SLC6A8 appear as a distinct metabolic feature of SqCC. Considering ongoing clinical trials focused on SLC6A8 inhibition in other malignancies, exploring SLC6A8 inhibition in SqCC appears motivated based on a metabolic addiction hypothesis.

## INTRODUCTION

Lung cancer is the leading cause of cancer-related mortality worldwide, with an estimated 1.8 million deaths in 2020. ^1^ The disease can broadly be divided into different histological subgroups, including adenocarcinoma (AC), squamous cell carcinoma (SqCC), large cell carcinoma (LCC), large cell neuroendocrine carcinoma (LCNEC), and small cell lung cancer (SCLC). In the current WHO guidelines, ^2^ LCNEC and SCLC form the neuroendocrine (NE) subgroup and the remaining histological subgroups form the non-small cell lung cancer (NSCLC) subgroup.

Extensive genomic profiling has identified a growing number of actionable alterations suitable for inhibition by molecular agents in lung cancer. The most successful applications are found in AC (the most common histological subgroup) and involve the targeting of mutation or fusion gene-activated protooncogenes like *EGFR*, *ALK*, *ROS1*, and *BRAF* for which molecular diagnostics are now clinical routine. ^3,4^ However, these activating mutations are typically not common in other histological subgroups, and targeted agents developed against these alterations are therefore of lesser clinical value outside of AC. ^5^ Identification of other actionable alterations in e.g., both early and advanced stage SqCC (the second most common histological subgroup) are of high clinical value.

In parallel to extensive DNA, RNA, and protein characterization of lung cancer, metabolomic profiling of the disease has also been performed. Importantly, metabolites and their concentrations directly reflect the underlying biochemical activity and state of cells/tissues and altered tumor cell metabolism has been identified as a hallmark of carcinogenesis. ^6^ Changes in tumor metabolism involve deregulated metabolism of glucose and amino acids, opportunistic modes of nutrient acquisition, increased nucleotide and polyamine biosynthesis, alterations in metabolite-driven gene regulation, and metabolic interaction with the tumor microenvironment. ^6–8^ Metabolic-oriented studies in lung cancer have identified biomarkers for early diagnosis, predicted prognosis by comparing changes in metabolites before and after surgery, discovered possible metabolic pathways of the disease, and attempted to determine lung cancer staging. ^9–13^ Moreover, recent studies have confirmed that the AC and SqCC histological subgroups appear to be associated with metabolic differences. ^12–16^

The aim of the current study was to identify genes and pathways associated with metabolic differences specific to SqCC histology, which might serve as new alternatives for future therapeutic intervention. Based on the integration of matched transcriptional, proteogenomic, and metabolomic data, key genes differentially expressed in SqCC were mapped to metabolite changes through genome-scale metabolic models (GEMs) which is a network-based tool that collects all known metabolic information of a biological system to computationally describe gene-protein-reaction associations. We specifically identified the creatine metabolism as enriched in SqCC tumors, with associated elevated levels of the creatine transporter *SLC6A8* gene in SqCC tumor cells, indicating specific metabolic fingerprints of lung cancer histologies. Notably, therapeutic inhibition of *SLC6A8* is currently tested in other malignancies serving as one example of potential new future treatment options in SqCC.

## MATERIAL AND METHODS

### Patient cohorts

A multi-omics tumor discovery cohort consisting of 156 NSCLC patients from our previously reported cohort of 159 RNA, DNA, and DNA-methylation profiled early-stage lung cancers surgically treated at the Skåne University Hospital in Lund, Sweden was used to identify genes and metabolites associated with SqCC histology. ^17^ All patients had early-stage disease, except for two patients (LU1113 and LU1068) who were treated as operable cases and underwent surgical resection, but whose tumors were subsequently diagnosed as stage IV. 73/156 cases had sufficient amounts of fresh-frozen tumor tissue for metabolic analysis. Histological subgroup classification was performed by a pathologist (H.B.) according to the WHO 2015 classification. ^18^ Patient characteristics are summarized in Table 1. In addition to the discovery cohort, we collected a cohort of 27 patients diagnosed with stage IV lung cancer at the Skåne University Hospital in Lund, Sweden, with tissue material collected through bronchoscopy at the time of diagnosis. Patient characteristics are summarized in Table 2 and this cohort is referred to as the Advanced LUCAS cohort.

**Table 1.** Clinicopathological characteristics of the tumor discovery cohort.

|  | All cases | AC | SqCC | LCC | LCNEC |
| --- | --- | --- | --- | --- | --- |
| **Nbr (%)** | 156 (100%) | 102 (65.4%) | 30 (19.2%) | 10 (6.4%) | 14 (9.0%) |
| **Gene expression data** | 156 (100%) | 102 (100%) | 30 (100%) | 10 (100%) | 14 (100%) |
| **Proteogenomic data** | 118 (75.6%) | 79 (77.5%) | 25 (83.3%) | 4 (40%) | 10 (71.4%) |
| **Metabolomics data** | 73 (46.8%) | 37 (36.3%) | 17 (56.7%) | 9 (90%) | 10 (71.4%) |
| **Median age (range)** | 67.9 (33.9-84) | 67 (36-83) | 72 (59-84) | 64.5 (47.6-76.2) | 61.1 (33.9-76.9) |
| **Gender (%)** |  |  |  |  |  |
| Female | 82 (52.6%) | 59 (57.8%) | 11 (36.7%) | 3 (30%) | 9 (64.3%) |
| Male | 73 (46.8%) | 43 (42.2%) | 18 (60%) | 7 (70%) | 5 (35.7%) |
| NA | 1 (0.6%) | 0 (0%) | 1 (3.3%) | 0 (0%) | 0 (0%) |
| **Smoking status** |  |  |  |  |  |
| Never-smoker | 19 (12.2%) | 18 (17.6%) | 1 (3.3%) | 0 (0%) | 0 (0%) |
| Smoker | 111 (71.2%) | 71 (69.7%) | 25 (83.3%) | 5 (50%) | 10 (71.4%) |
| NA | 26 (16.6%) | 13 (12.7%) | 4 (13.4%) | 5 (50%) | 4 (28.6%) |
| **Stage** |  |  |  |  |  |
| I | 119 (76.3%) | 82 (80.4%) | 26 (86.7%) | 3 (30%) | 8 (57.2%) |
| II | 26 (16.7%) | 13 (12.7%) | 3 (10%) | 5 (50%) | 5 (35.7%) |
| III | 7 (4.5%) | 5 (4.9%) | 0 (0%) | 1 (10%) | 1 (7.1%) |
| IV | 2 (1.3%)** | 0 (0%) | 1 (3.3%)** | 1 (10%)** | 0 (0%) |
| NA | 2 (1.3%) | 2 (2%) | 0 (0%) | 0 (0%) | 0 (0%) |
| ***EGFR* mutation \*** | 10 (7.4%) | 10 (10%) | 0 (0%) | 0 (0%) | 0 (0%) |
| ***KRAS* mutation\*** | 31 (23%) | 31 (31%) | 0 (0%) | 0 (0%) | 0 (0%) |

\* percent of patients tested that were positive for the mutation
\*\*Patients LU1113 and LU1068 were treated as operable cases and underwent surgical resection, but the tumors were subsequently diagnosed as stage IV tumors.

**Table 2.** Clinicopathological characteristics of the advanced LUCAS cohort.

|  | All cases | AC | SqCC | NSCLC- NOS | SCLC |
| --- | --- | --- | --- | --- | --- |
| **Nbr (%)** | 27 (100%) | 15 (55.6%) | 6 (22.2%) | 3 (11.1%) | 3 (11.1%) |
| **Median age (range)** | 75 (46-95) | 77 (55-95) | 71 (62-80) | 63 (46-72) | 78 (77-78) |
| **Gender (%)** |  |  |  |  |  |
| Female | 14 (52%) | 9 (60%) | 3 (50%) | 1 (33%) | 1 (33%) |
| Male | 13 (48%) | 6 (40%) | 3 (50%) | 2 (67%) | 2 (67%) |
| **Smoking status** |  |  |  |  |  |
| Current | 7 (26.9%) | 5 (35.7%) | 1 (16.7%) | 0 (0%) | 1 (33.3%) |
| Former | 17 (65.4%) | 7 (50%) | 5 (83.3%) | 3 (100%) | 2 (66.7%) |
| Never-smoker | 2 (7.7%) | 2 (14.3%) | 0 (0%) | 0 (0%) | 0 (0%) |
| **Stage** |  |  |  |  |  |
| IV A | 14 (51.9%) | 8 (53.3%) | 5 (83.3%) | 1 (33.3%) | 0 (0%) |
| IV B | 13 (48.1%) | 7 (46.7%) | 1 (16.7%) | 2 (66.7%) | 3 (100%) |
| **EGFR mutation** | 3 (11%) | 3 (20%) | 0 | 0 | 0 |
| **ALK fusion** | 1 (4%) | 1 (7%) | 0 | 0 | 0 |

### Gene expression and proteogenomic data for discovery and Advanced LUCAS cohorts

Processed gene expression data for cases in the tumor discovery cohort was obtained from Gene Expression Omnibus (GSE94601). ^17^ Processed proteogenomic data for 118 of the 156 cases was obtained from Lethiö et al. ^19^

For the Advanced LUCAS cohort, RNA was extracted from 27 bronchoscopy samples (tissue pieces or fine needle aspirations stored in RNAlater solution (Invitrogen) directly at the time of clinical investigation), using the Allprep DNA/RNA mini kit (Qiagen). RNA sequencing (RNA-seq) was performed at the Center for Translational Genomics (CTG) at Lund University using the Illumina Stranded mRNA library preparation protocol and sequenced on a Novaseq instrument (Illumina). Data processing was performed as previously described to generate fragments per kilobase of transcript per million reads (FPKM) expression estimates.^20^

### Public gene expression and metabolite data

Paired tumor and normal RNA-seq data from 57 patients with AC and 49 patients with SqCC tumors were obtained from The Cancer Genome Atlas (TCGA). Data was downloaded from the National Cancer Institute’s Genomic Data Commons Data Portal (portal.gdc.cancer.gov) in the form of GDS RNAseqv2 FPKM. FPKM data was upper quantile normalized (“FPKM-UQ”), followed by log2 transformation using a +1 offset prior to log2 transformation.

Tumor gene expression data for 183 lung tumors was obtained from Djureinovic et al., ^21^ including 115 AC and 68 SqCC. Gene expression data for 19144 genes for 185 lung cancer cell lines was obtained from the DepMap portal (www.depmap.org, accessed October 29, 2020). Based on subgroup annotations provided by the repository, the cell lines were divided into 79 AC, 27 SqCC, 10 NSCLC not otherwise specified (NSCLC-NOS), 17 LCC, and 52 SCLC. From the same repository, we also obtained matched normalized proteomic data for 12755 proteins for 75 cell lines (37 AC, 12 SqCC, 4 NSCLC-NOS, 9 LCC, and 13 SCLC). Corresponding metabolite expression for 225 metabolites for 168 cell lines (69 AC, 23 SqCC, 8 NSCLC-NOS, 17 LCC, and 51 SCLC) was obtained from Li et al. ^22^

### SqCC specific genes

SqCC-specific genes were derived through differential gene expression performed on median-centered expression data in the discovery cohort. Multigroup comparison was performed by Kruskal-Wallis testing and genes with Benjamini-Hochberg adjusted p-value <0.05 were retained. Paired Wilcoxon tests were then performed for each of these genes and for each subtype combination. Genes that had exactly three tests with Benjamini-Hochberg adjusted p-value <0.05, all connected to the SqCC histology, were retained as differentially expressed in SqCC only. Each of the SqCC-specific genes were then tested for significance in the Djureinovic cohort. ^21^

### Metabolomics

For 73 tumors from the discovery cohort that had sufficient amounts of fresh-frozen tumor tissue available, 20 to 30 mg of tissue was pulverized via mechanical disruption (TissueLyser II, Qiagen) prior to hydrophilic extraction of intracellular metabolites from tissue using a methanol/acetonitrile/water (50/30/20) with 5 µM d8-valine extraction solution (250 μL of extraction solution per 10mg homogenized tissue). Following a one-hour incubation at -20°C, the samples were placed in a Thermomixer for 15 minutes at 4°C and maximum speed. The samples were then centrifuged for 10 minutes at maximum speed and the supernatant was kept at -80°C in autosampler vials prior to mass spectrometry analysis.

HILIC chromatographic separation of metabolites was achieved using a Millipore Sequant ZIC-pHILIC analytical column (5 µm, 2.1 × 150 mm) equipped with a 2.1 × 20 mm guard column (both 5 mm particle size) with a binary solvent system. Solvent A was 20 mM ammonium carbonate, 0.05% ammonium hydroxide; Solvent B was acetonitrile. The column oven and autosampler tray were held at 40°C and 4°C, respectively. The chromatographic gradient was run at a flow rate of 0.200 mL/min as follows: 0–2 min: 80% B; 2-17 min: linear gradient from 80% B to 20% B; 17-17.1 min: linear gradient from 20% B to 80% B; 17.1-22.5 min: hold at 80% B. Samples were randomized and analyzed with Liquid chromatography– mass spectrometry (LC–MS) in a blinded manner with an injection volume 5 µl. Pooled samples were generated from an equal mixture of all individual samples and analyzed interspersed at regular intervals within sample sequence as a quality control.

Metabolites were measured with a Thermo Scientific Q Exactive Hybrid Quadrupole-Orbitrap Mass spectrometer (HRMS) coupled to a Dionex Ultimate 3000 UHPLC. The mass spectrometer was operated in full-scan, polarity-switching mode, with the spray voltage set to +4.5 kV/-3.5 kV, the heated capillary held at 320 °C, and the auxiliary gas heater held at 280°C. The sheath gas flow was set to 25 units, the auxiliary gas flow was set to 15 units, and the sweep gas flow was set to 0 units. HRMS data acquisition was performed in a range of m/z = 70–900, with the resolution set at 70,000, the AGC target at 1 × 106, and the maximum injection time (Max IT) at 120 ms. Metabolite identities were confirmed using two parameters: (1) precursor ion m/z was matched within 5 ppm of theoretical mass predicted by the chemical formula; (2) the retention time of metabolites was within 5% of the retention time of a purified standard run with the same chromatographic method. Chromatogram review and peak area integration were performed using the Thermo Fisher software Tracefinder 5.0 and the peak area for each detected metabolite was normalized against the total ion count (TIC) of that sample to correct any variations introduced from sample handling through instrument analysis. The normalized areas were used as variables for further statistical data analysis. TIC-normalized data for 139 metabolites and 73 cases was used in subsequent analyses.

### Immunohistochemical evaluation of SLC6A8 expression

Immunohistochemical (IHC) staining was performed on a prospective cohort including 213 patients with primary lung cancer who underwent surgical treatment at the Skåne University Hospital, Lund between 2005 and 2011. The study included 134 AC, 11 NE tumors, and 68 SqCC cases and has been described previously. ^23–25^ In this cohort, one LCNEC case, 18 AC, and 12 SqCC cases overlap with the tumor discovery cohort.

Tissue microarrays (TMA) were used for IHC analysis. The TMA blocks had three cores of 1 mm in diameter per sample (for rare cases, tumor cells were present on only one or two of the cores). For IHC analysis, 4 µm thick sections were stained for SLC6A8 on a Ventana Benchmark automated staining platform using a rabbit polyclonal antibody (Proteintech, Germany) at a 1:200 dilution. The slides were scanned and evaluated using the pathXL software (Philips, The Netherlands). Stained slides were evaluated by two independent observers (D.E. and M.J.) who were blinded to clinical data and patient outcomes; for cases with differences in the scoring between evaluators, a third observer (H.B.) was consulted, and consensus was reached. In all evaluated cases there were at least 200 tumor cells, and in the clear majority of cases, more than 1000 evaluable cells. Complete membrane IHC staining of tumor cells, or partial membrane staining when discernable to a single cell, was considered positive. Tumor cells with exclusively cytoplasmic staining were disregarded. Special care was taken to only count positive tumor cells, and to exclude other protein-expressing cells, such as macrophages or respiratory epithelial basal cells. Complementary IHC such as CD68 for macrophages as well as hematoxylin-eosin sections of the cores were available for all cases when needed. Protein expression of SLC6A8 in tumor cells was scored using a four-graded scale: <1%, 1-10%, 11-50%, or >50%.

### Statistics

P-value adjustment was performed using the p.adjust function and functional pathway analysis of biological processes was performed using the enrichGO function, both in R (www.r-project.org). Survival analysis was performed using the online KMplotter tool, ^26^ with overall survival (OS) as the clinical endpoint. In KMplotter, SqCC tumors were divided into three equally sized groups based on mRNA levels for investigated genes and univariate Cox regression using the lowest expressing group versus the highest expressing group was performed to assess prognostic association. The online MetaboAnalyst version 4.0 tool ^27^ was used for pathway analysis of metabolite networks using default settings and all compounds in the selected pathways as reference sets. The R piano package version 2.14.0 was used to map metabolites to the genes of interest through genome-scale metabolic models.

### Ethics statement

The study was approved by the Regional Ethical Review Board in Lund, Sweden (Registration no. 2004/762, 2008/702, and 2014/748) and performed in adherence with the Declaration of Helsinki.

## RESULTS

### SqCC specific gene expression to identify potential metabolic associations

An overview of the study is provided in Figure 1. Differential gene expression analysis in the discovery cohort identified 280 genes with mRNA expression specific to SqCC (Supplementary Table S1). Of the 280 SqCC-specific genes, 212 were present in the RNA-seq bulk data for 183 early-stage lung tumors from Djureinovic et al. ^21^ and of these, 194 genes were also significantly differentially expressed between AC and SqCC. Functional pathway analysis of the 280 genes identified 66 enriched biological processes, of which several were expected considering upregulation in SqCC of numerous keratin genes like *KRT6A*, *KRT17*, *KRT16*, and *KRT15*, including tissue development (GO:0009888), epidermis development (GO:0008544), epithelial cell differentiation (GO:0030855), and keratinocyte differentiation (GO:0030216) (Supplementary Table S1). No apparent associations with metabolic pathways were identified, despite reported metabolic differences between histological lung cancer subtypes. ^12,13^ The latter may be due to only a single/few key genes in a metabolic pathway, e.g., a transporter protein, showing large fold-changes in global differential gene expression analysis. To more comprehensively search for metabolic associations of the 280 genes we used Genome-scale metabolic models (GEMs) from the Metabolic Atlas to identify metabolites whose pathway involved at least one differentially expressed gene. Notably, 57 of the 280 SqCC-specific genes were connected to 324 unique metabolites through GEMs (Supplementary Table S1). The top-ranking differentially expressed genes associated with a metabolite included, e.g., the creatine transporter gene *SLC6A8* associated to creatine, the interferon Regulatory Factor 6 gene *IRF6* associated to phenylalanine, the SH3 Domain Binding Protein 1 gene *SH3BP1* associated to pyridoxal, and the cyclin Dependent Kinase 6 gene *CDK6* associated to ATP.

**Figure 1:**
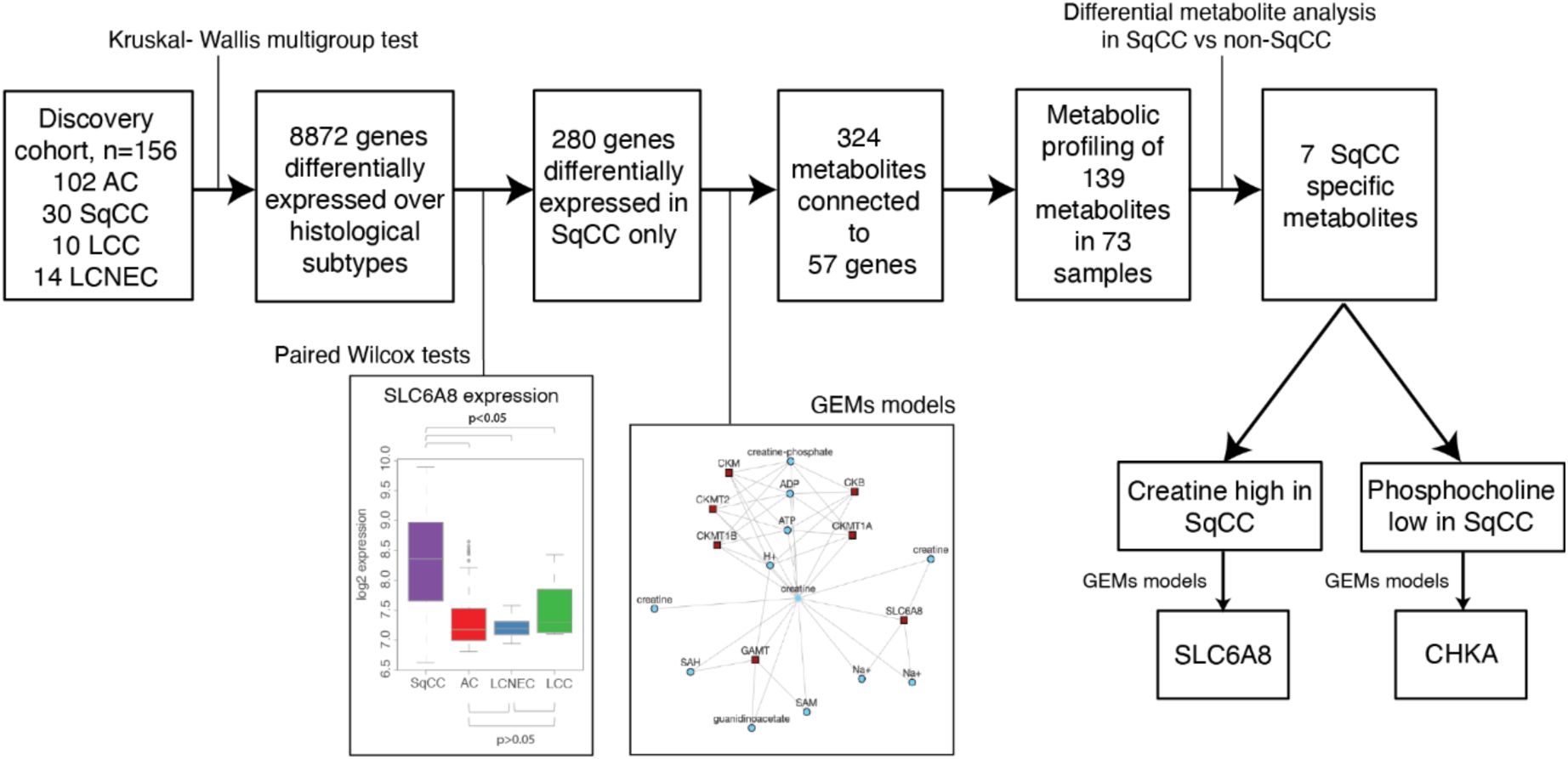
Overview of study with key findings.

### Metabolic profiling of lung cancers identifies two metabolites specific to SqCC

To investigate the connection between the identified SqCC-specific gene and relative metabolic patterns in early-stage lung cancer we performed mass spectrometry-based metabolic profiling of 73 tumors from the tumor discovery cohort targeting 139 metabolites (Supplementary Table S2). Of the 139 measurable metabolites, 24 could be mapped through GEMs to 34 genes from the 280 gene list (Supplementary Table S1). Based on a one-vs-rest approach (SqCC vs non-SqCC), seven metabolites (creatine, guanine, guanosine, N-acetylneuramic acid, phosphocholine, xanthine, and homocitrulline) were identified as having significantly different levels in SqCC versus non-SqCC tumors (Figure 2A).

**Figure 2.**
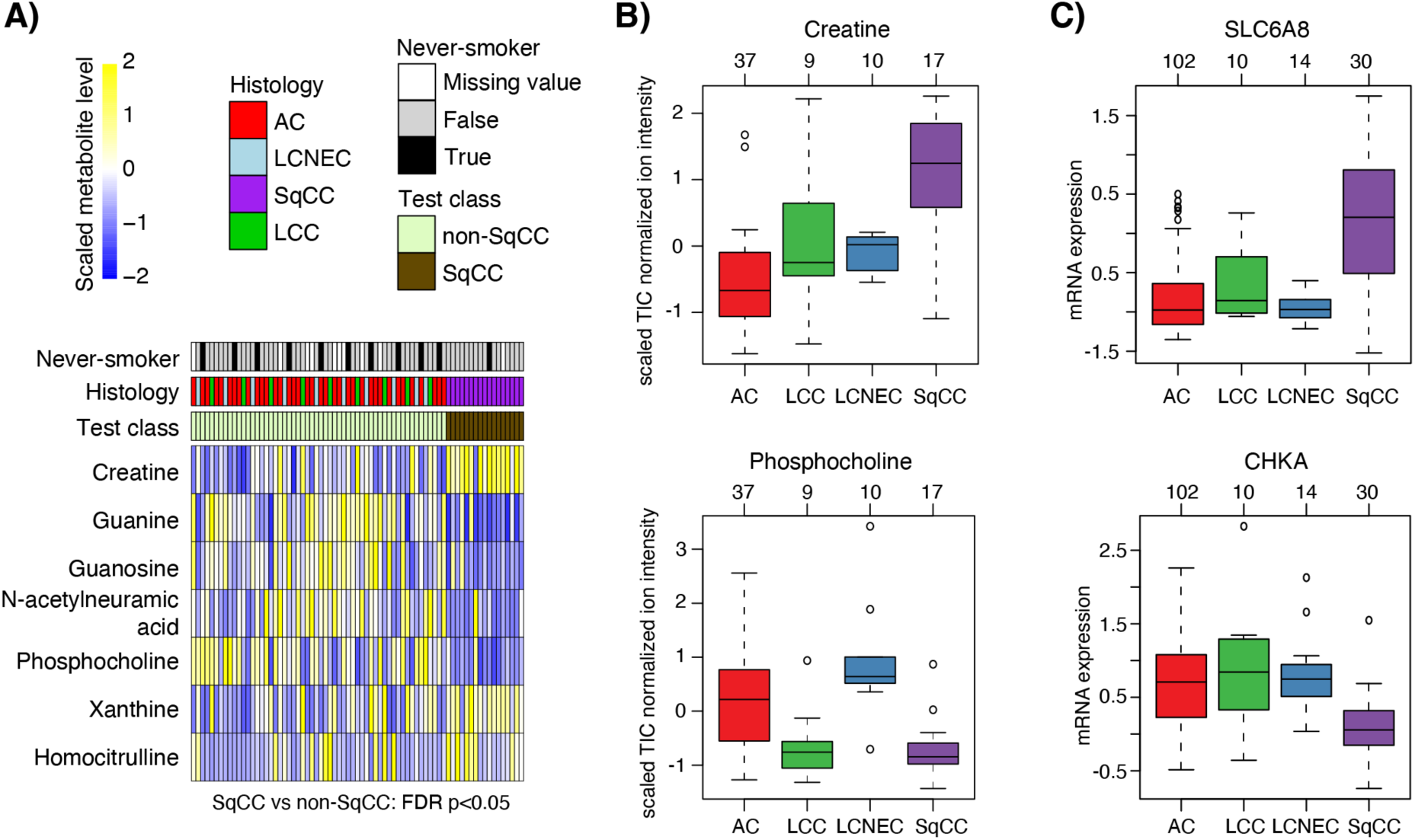
SqCC-specific metabolites and matching gene expression in lung cancer cases from the discovery cohort. (A) Ordered heatmap of scaled metabolite data for metabolites found to be different in abundance between SqCC and non-SqCC cases in 73 tumors (FDR adjusted two-sided Wilcoxon’s p-value <0.05). (**B)** Scaled creatine and phosphocholine metabolite levels versus histological subtypes. (**C)** mRNA expression of *SLC6A8* and *CHKA* associated with creatine and phosphocholine, respectively, across the histological subtypes.

Of the seven significant metabolites, two -creatine (elevated in SqCC) and phosphocholine (decreased in SqCC)- involved genes (the creatine transporter gene *SLC6A8* and the choline kinase A gene *CHKA*) were found to have SqCC-specific mRNA expression (Supplementary Table S1). Notably, the mRNA expression of both *SLC6A8* and *CHKA* matched the corresponding metabolite level in the tumor tissue (Figure 2C).

### Targeting SLC6A8 and CHKA associated with SqCC metabolite-specific expression

To substantiate the expression patterns of *SLC6A8* and *CHKA* in lung cancer we first compared the mRNA expression of the two genes in the Djureinovic et al. RNA-seq cohort, finding gene expression patterns of both genes closely matching those observed in our discovery cohort (Figure 3A). Next, we evaluated the gene expression of *SLC6A8* and *CHKA* in matched tumor/normal data for 57 AC and 49 SqCC cases from the TCGA consortium. Again, similar gene expression patterns were observed across the histologies in the TCGA cases compared to our tumor discovery cohort, providing further validation of the latter. Moreover, the mRNA levels of these key genes appeared altered in tumor tissue compared to expression in matched normal tissue, suggesting tumor-specific expression (Figure 3B). To analyze the mRNA expression of the two genes also in advanced lung cancer, we performed RNA-seq on freshly collected bronchoscopy material (tissue and fine needle aspirations, as outlined ^28^) at the time of clinical investigation from 27 stage IV patients (15 AC, 6 SqCC, 3 SCLC, and 3 NSCLC-NOS) (Supplementary Table S3). Consistent with the discovery cohort we observed elevated *SLC6A8* mRNA expression in stage IV SqCC compared to non-SqCC tumors, while *CHKA* was not significantly expressed (Figure 3C).

**Figure 3.**
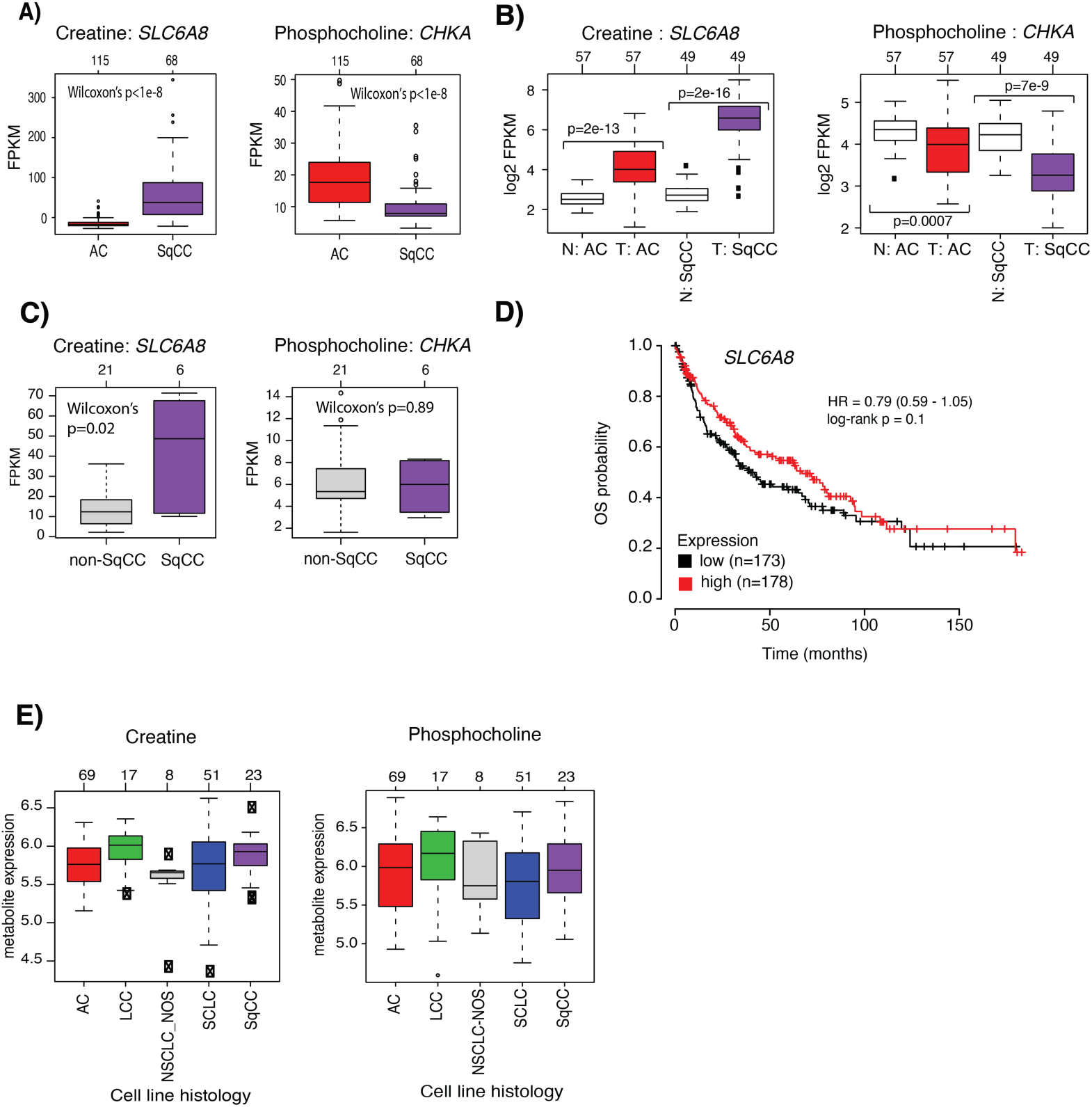
Metabolite abundance of creatine and phosphocholine and associated genes in tumor, matched tumor-normal tissue, and cancer cell lines. (**A**) Gene expression of the *SLC6A8* and *CHKA* genes associated with the metabolites creatine and phosphocholine, respectively, in SqCC versus AC tumors from Djureinovic et al. ^21^ P-values were calculated using a two-sided Wilcoxon’s test. (**B**) Gene expression of *SLC6A8* and *CHKA* in matched tumor to normal tissue from 57 patients with AC and 49 patients with SqCC from the TCGA consortium. ‘N AC’: matched normal tissue for patients with AC tumors. ‘N SqCC’: matched normal tissue for patients with SqCC tumors. (**C)** Gene expression of *SLC6A8* and *CHKA* for SqCC versus non-SqCC tumor histologies in 27 stage IV lung tumors. (**D)** Kaplan-Meier plot with OS as the clinical endpoint for patients grouped based on *SLC6A8* gene expression divided into two equally sized groups based on mRNA expression. Analysis was performed using the online KMplotter tool. (**E)** Creatine and phosphocholine metabolite concentrations versus histological subtype in lung cancer cell lines. Metabolite levels were obtained from the study by Li et al. ^22^

Finally, we tested the prognostic value of *SLC6A8* in 524 primary SqCC tumors using the online KMplotter tool ^26^, dividing cases into three equally sized groups based on mRNA expression. Based on univariate Cox regression, no significant association with OS was observed for the group with the lowest versus highest mRNA levels (hazard ratio=0.79, 95% confidence interval=0.59-1.05; Figure 3D). Using the same tool, we found similar non-significant results for *CHKA* (log-rank p>0.05).

### mRNA, protein, and matched metabolite expression of SqCC-specific SLC6A8 and CHKA in public cancer cell line data

To further investigate SLC6A8 and CHKA in lung cancer we analyzed mRNA, protein, and metabolite expression of the two genes in lung cancer cell lines. For mRNA and protein patterns we collected 185 lung cancer cell lines divided into 79 AC, 27 SqCC, 10 NSCLC-NOS, 17 LCC, and 52 SCLC cell lines from the DepMap consortium. However, gene consistency across histological subgroups of lung cancer cell lines was poor for mRNA and protein expression when compared to the tumor data. *SLC6A8* showed a consistent protein, but not mRNA expression pattern, compared to the tumor tissue data, while CHKA did not (Supplementary Figure S1).

Li et al. recently reported metabolomic profiling of a large number of cancer cell lines from different organs. ^22^ In this cohort, we identified 168 lung cancer cell lines divided into 69 AC, 23 SqCC, 8 NSCLC-NOS, 17 LCC, and 51 SCLC with data for 225 metabolites. However, we did not observe the same patterns of creatine and phosphocholine concentrations as in tumor tissue (Figure 3E).

### SLC6A8 protein expression in early-stage lung cancer

To substantiate protein expression patterns of SLC6A8 in the discovery cohort, we investigated the expression of the gene using IHC in a TMA comprising 213 lung cancers (134 AC, 11 NE, and 68 SqCC tumors). ^23–25^ Protein expression was scored either on a four-graded scale: <1%, 1-10%, 11-50%, or >50% or as a binary variable (<1% or ≥1%) for the mean IHC score across TMA cores. Consistent with mRNA and metabolite expression patterns in previous cohorts, SqCC tumors showed significantly higher SLC6A8 protein expression than AC and NE tumors (Figure 4, Chi-square test p<2e-16 for all comparisons). Pathologist (D.E., H.B.) evaluation of protein expression established that the protein was primarily expressed in tumor cells. Respiratory basal cells were also expressors, as were some macrophages, although the latter mainly exhibited vague cytoplasmic staining and not the membranous staining that was considered positive. Necrotic debris could manifest with some slight positivity in IHC, but these were mostly easy to disregard.

**Figure 4.**
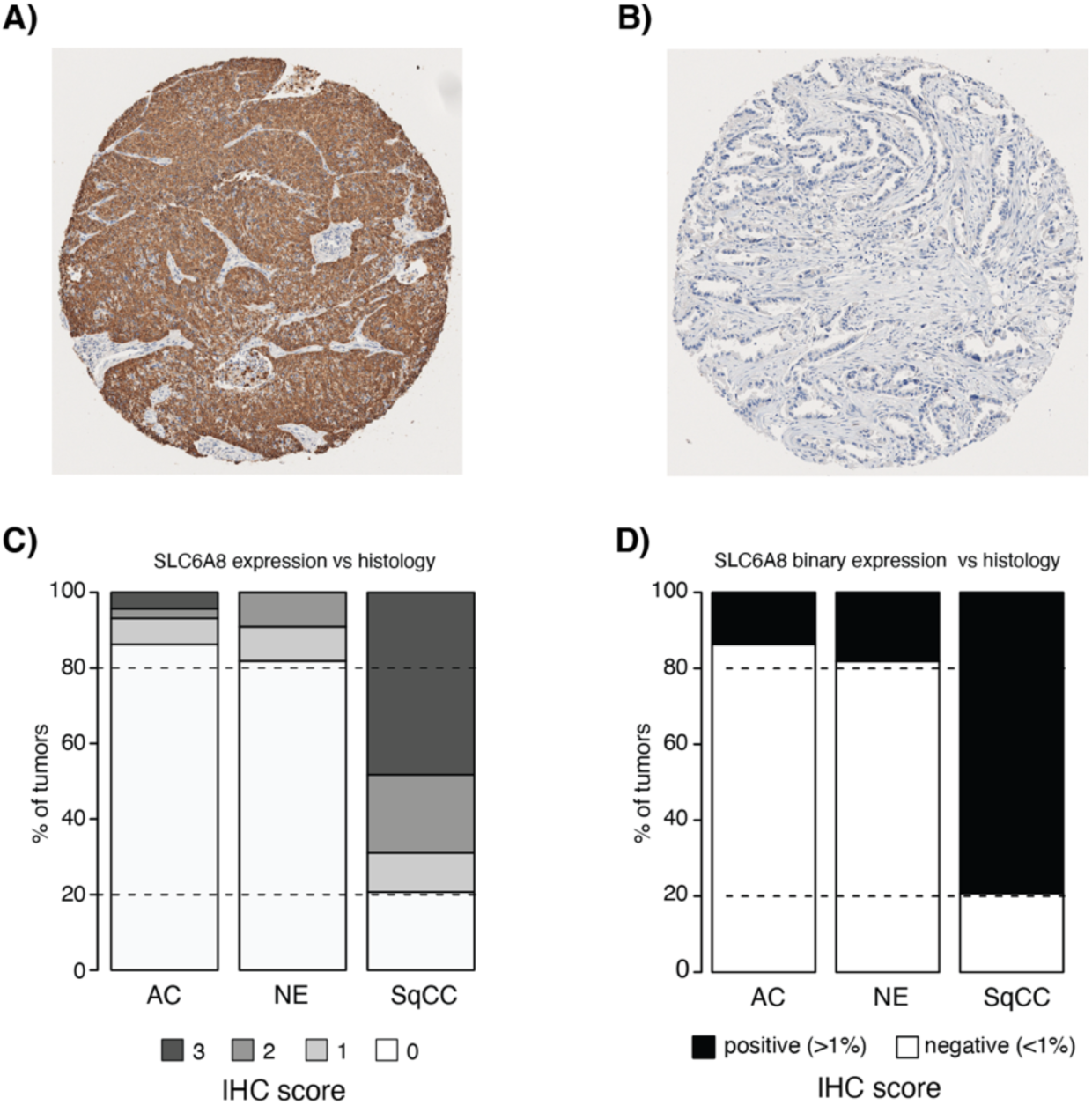
SLC6A8 protein expression versus histological subtypes. (**A**) SLC6A8 IHC stain of a SqCC tumor with assessed >50% mean staining. (**B**) SLC6A8 IHC stain of an AC tumor with assessed <1% mean staining. (**C**) SLC6A8 IHC scores (0=<1%,1=1-10%,2=10-50%, 3= >50%) versus histological subtype. (**D**) SLC6A8 IHC binary scores (negative <1%, positive ≥1%) versus histological subtype.

## DISCUSSION

Metabolic alterations are associated with several pathological conditions, including cancer. Metabolomic profiling of malignancy may thus be a powerful approach to complement findings from other -omics technologies to improve disease understanding and any molecular phenotypes within. In this study, we searched for metabolite-connected genes specific to SqCC by integrating transcriptomics, proteomics, and mass spectrometry-based metabolomic analysis in both tumor tissue and cancer cell lines. In tumor tissue, this approach identified two metabolites, creatine and phosphocholine, specifically associated with SqCC histology with matched gene and protein expression of key genes included in corresponding metabolic models.

Deregulated creatine (in particular) and phosphocholine metabolite levels between AC and SqCC have been reported previously, ^12,13,29,30^ with similar patterns as observed in this study, but without the link to SLC6A8 or CHKA expression through metabolic models. When comparing bulk tumor tissue to matched normal tissue, the increased mRNA expression of the creatine transporter *SLC6A8* (solute carrier family 6, member 8) gene and decreased *CHKA* (phosphocholine) expression appeared tumor-associated. The latter was further corroborated for SLC6A8 by IHC staining which showed that in situ expression of the SLC6A8 protein was mainly specific to tumor cells (Figure 4). These findings are in line with previous reports of high SLC6A8 protein expression in AC and NSCLC in general versus non-malignant tissue. ^31,32^ In the study by Feng et al. it was proposed that SLC6A8 overexpression promotes the proliferation, migration, and invasion in vitro in NSCLC, accompanied by the activation of notch signaling pathway. ^32^ In contrast, in human tumor tissue our results indicate that SLC6A8 overexpression is a distinctive feature of SqCC tumor cells that it is highly correlated with increased creatine levels in SqCC tumor tissue compared to other histological subtypes (suggesting a potential difference in tumor metabolism), but, importantly, that these observations replicate poorly in cancer cell line models.

The creatine transporter protein SLC6A8 imports the high-energy metabolite phosphocreatine into the cell where it can be converted to ATP to fuel the survival of cancer cells as they proliferate and spread. *SLC6A8* is overexpressed in several cancer types, including gastrointestinal cancers where creatine metabolism has been implicated in colon cancer progression and metastatic colonization of the liver. ^33,34^ Here, cancer cells may upregulate and release creatine kinase-B (CKB) into the extracellular space where it generates the high-energy metabolite phosphocreatine that is transported into the cell by the SLC6A8 transporter protein. Consistent with this finding, genetic depletion of SLC6A8 in colon cancer cell lines significantly reduced cancer growth in animal studies. ^33^ In ongoing clinical phase 1 and 2 trials, an oral small molecule inhibitor (RGX-202) of SLC6A8 is tested as a single agent or as combination therapy in patients with advanced gastrointestinal malignancies (ClinicalTrials.gov ID NCT03597581 and NCT05983367, and Kurth et al. ^34^). If SLC6A8 inhibition can reduce intracellular levels of phosphocreatine available for ATP synthesis in tumor cells, tumor cell growth and metastasis may potentially be limited. Based on our findings of consistently high creatine levels in SqCC tumor cells compared to other histological subgroups, it appears imminent to also investigate the potential of SLC6A8 inhibitors in SqCC treatment. This brings forward an intriguing hypothesis of potential metabolite addiction in SqCC, as a parallel to the concept of oncogene addiction in AC based on recurrent oncogenic alterations in different tyrosine kinases.

To ensure reproducibility, we consistently compared our results to different public tumor cohorts and tested our findings also in cancer cell line data. A notable finding was similar patterns of gene expression and/or protein expression in the different tumor cohorts, but not in cancer cell lines. Specifically, congruence between metabolite, mRNA, and protein levels for cell lines as models of SqCC and actual measurements in tumor tissue appeared to be modest, especially for measured metabolite concentrations, an observation reported also by others for 2D cultures of established cell lines. ^14^ The lack of agreement may be due to different metabolite analysis methods but could also be related to the nature and culture conditions of immortalized human cancer cell lines. Here, the additional validation of key genes in external tumor and normal tissue data provides greater support to the original tumor discovery cohort findings compared to the less supportive cell line data, while also highlighting a potential problem with cell lines as representative in vitro models for metabolic changes in tumors (in line with ^14^).

Limitations of the current study include the lack of available matched normal tissue in the discovery cohort for matched metabolic profiling. Moreover, the metabolic profiling in the current study is focused on hydrophilic metabolites, thus generating a small set of measured metabolites across all tumors. Moreover, in lung cancer (including SqCC) molecular phenotypes based on gene expression, proteomics, epigenetics, and genetic alterations have been reported. ^5,19,35–37^ It is plausible, but not yet proven, that such molecular subtypes may also involve changes in cell metabolism within the histological subgroups. Due to the limited size of the current study and the unavailability of matched normal controls, we were not able to account for proposed/potential molecular subtypes in the identification of histology-associated metabolites. However, for the key finding of the creatine / SLC6A8 connection, the unified high expression in SqCC tumors would argue against large differences within SqCC due to specific molecular phenotypes.

The clinical need for targeted therapy in SqCC is urgent. Current targeted therapies in lung cancer are directed towards genetic alterations in oncogenes rarely found in SqCC tumors^3^, leaving advanced-stage patients the options of immunotherapy and/or chemotherapy. In this study, we demonstrate that the SLC6A8 creatine transporter gene is overexpressed in both early and advanced stage SqCC. The generally high levels of creatine in SqCC suggest that it is not a prognostic variable within the subgroup, but rather a subgroup-specific alteration that may be actionable. The most intuitive method to test SLC6A8 inhibition in SqCC would be using cell lines as in vitro models. However, our analyses of public cell line data indicate that cell lines are not the optimal model as there is discordance in metabolite levels and mRNA and protein expression of *SLC6A8* compared to tumor tissue results. Therefore, alternative in vitro or in vivo models would be needed to test the potential of SLC6A8 inhibition as a targeted SqCC treatment. A main question to answer would be how strong the metabolic dependency is during treatment pressure, i.e., whether tumor cells can easily adapt and change their metabolism to overcome inhibition.

In summary, based on integrated analysis of high-dimensional transcriptomic and proteogenomic data, with metabolite profiling we identified two metabolites in SqCC with associated gene and/or protein expression of key metabolite network genes, linking previous findings from studies in the field into a coherent metabolite-gene framework and raising a speculation about a potential metabolic addiction in SqCC to be explored in vivo. Interestingly, the elevated levels of creatine and its transporter protein SLC6A8 may be a potential drug target in SqCC based on oral inhibitors already in clinical phase 1 and 2 use in gastrointestinal malignancies.

## ADDITIONAL INFORMATION

### Acknowledgments

The authors would like to acknowledge Clinical Genomics Lund, SciLifeLab, and Center for Translational Genomics (CTG), Lund University, for providing expertise and service with sequencing and analysis.

### Authors’ contributions

EA, JS, MP and CF conceptualised the study. EA, AK, MJ and FR carried out experiments. CF, DE, HB, MK and MP provided clinical expertise and/or resources. MA, JS, CF and EA carried out data curation. EA and JS analysed the data. EA, JS, CF, MP and DE acquired funding. All authors were involved in writing the paper and had final approval of the submitted and published version.

### Ethics approval and consent to participate

The study was approved by the Regional Ethical Review Board in Lund, Sweden (Registration no. 2004/762, 2008/702, and 2014/748).

The study was performed in adherence with the Declaration of Helsinki.

### Consent for publication

Not applicable

### Data availability

Metabolomic data for the LU discovery cohort is available in Supplementary table S2. RNA-seq FPKM counts are available in Supplementary table S3.

Data from The Cancer Genome Atlas initiative can be retrieved through the Genomic Data Commons Data Portal at https://portal.gdc.cancer.gov.

Gene expression and proteomic data for lung cancer cell lines was obtained from the DepMap portal (www.depmap.org)

Other data is available from the corresponding publications as cited in the methods sections.

### Competing interests

The authors declare no potential conflicts of interest.

### Funding information

Financial support for this study was provided by the Swedish Cancer Society, the Sjöberg Foundation, the Mrs Berta Kamprad Foundation (2021-13, 2022-38), the Gustav V Jubilee Foundation, the Craaford Foundation (20220983), the Royal Physiographic Society of Lund (42366), Magnus Bergvalls Foundation (2021-04321, 2022-287), The Thelma Zoégas Foundation for Medicinal Research, The Stig and Ragna Gorthon Foundation and The National Health Services (Region Skåne/ALF). C.F. is funded by the ERC (ERC819920), the Alexander von Humboldt Foundation in the framework of the Alexander von Humboldt Professorship endowed by the Federal Ministry of Education and Research, and CRUK Programme Foundation award C51061/A27453.

The funders had no role in study design, data collection and analysis, decision to publish, or preparation of the manuscript

---

**Supplementary Figure S1.** Expression of matched metabolite, gene, and protein expression of (**A**) creatine and (**B**) phosphocholine in lung cancer cell lines.

**Supplementary Table S1:** A. 280 SqCC specific genes in the LU discovery cohort B. Pathway enrichment analysis of biological processes for the 380 SqCC specific genes C. Metabolites (all metabolites from the human GEMs metabolites or 139 measured metabolites) matched to 280 SqCC specific genes through GEMs.

**Supplementary Table S2:** A. TIC normalized ion intensity of 139 measured metabolites in 73 lung cancer samples B. 225 measured metabolites in 168 lung cancer cell lines from Li et al. ^22^.

**Supplementary Table S3:** FPKM RNA-seq data for the Advanced LUCAS cohort.

## SUPPLEMENTAL DATA

Supplementary figure S1.pdf

Supplementary table S1.xlsx

Supplementary table S2.xlsx

Supplementary table S3.xlsx

## Notes

### Competing Interest Statement

The authors have declared no competing interest.

## REFERENCES

1 Sung, H., Ferlay, J., Siegel, R. L., Laversanne, M., Soerjomataram, I., Jemal, A. et al. Global Cancer Statistics 2020: GLOBOCAN Estimates of Incidence and Mortality Worldwide for 36 Cancers in 185 Countries. CA Cancer J Clin 71, 209–249 (2021).

2 The WHO Classification of Tumours Editorial Board: Thoracic Tumours. 5th edn. Lyon: IARC Press; 2021.

3 Popper, H. H., Ryska, A., Timar, J. & Olszewski, W. Molecular testing in lung cancer in the era of precision medicine. Transl Lung Cancer Res 3, 291–300 (2014).

4 Shea, M., Costa, D. B. & Rangachari, D. Management of advanced non-small cell lung cancers with known mutations or rearrangements: latest evidence and treatment approaches. Ther Adv Respir Dis 10, 113–129 (2016).

5 Cancer Genome Atlas Research, N. Comprehensive genomic characterization of squamous cell lung cancers. Nature 489, 519–525 (2012).

6 Hanahan, D. & Weinberg, R. A. Hallmarks of cancer: the next generation. Cell 144, 646–674 (2011).

7 DeBerardinis, R. J. & Chandel, N. S. Fundamentals of cancer metabolism. Sci Adv 2, e1600200 (2016).

8 Pavlova, N. N. & Thompson, C. B. The Emerging Hallmarks of Cancer Metabolism. Cell Metab 23, 27–47 (2016).

9 Deja, S., Porebska, I., Kowal, A., Zabek, A., Barg, W., Pawelczyk, K. et al. Metabolomics provide new insights on lung cancer staging and discrimination from chronic obstructive pulmonary disease. J Pharm Biomed Anal 100, 369–380 (2014).

10 Fan, T. W., Lane, A. N., Higashi, R. M., Farag, M. A., Gao, H., Bousamra, M. et al. Altered regulation of metabolic pathways in human lung cancer discerned by (13)C stable isotope-resolved metabolomics (SIRM). Mol Cancer 8, 41 (2009).

11 Robles, A. I. & Harris, C. C. Integration of multiple “OMIC” biomarkers: A precision medicine strategy for lung cancer. Lung Cancer 107, 50–58 (2017).

12 You, L., Fan, Y., Liu, X., Shao, S., Guo, L., Noreldeen, H. A. A. et al. Liquid Chromatography-Mass Spectrometry-Based Tissue Metabolic Profiling Reveals Major Metabolic Pathway Alterations and Potential Biomarkers of Lung Cancer. J Proteome Res 19, 3750–3760 (2020).

13 Rocha, C. M., Barros, A. S., Goodfellow, B. J., Carreira, I. M., Gomes, A., Sousa, V. et al. NMR metabolomics of human lung tumours reveals distinct metabolic signatures for adenocarcinoma and squamous cell carcinoma. Carcinogenesis 36, 68–75 (2015).

14 Sellers, K., Allen, T. D., Bousamra, M., 2nd, Tan, J., Mendez-Lucas, A., Lin, W. et al. Metabolic reprogramming and Notch activity distinguish between non-small cell lung cancer subtypes. Br J Cancer 121, 51–64 (2019).

15 Hoang, L. T., Domingo-Sabugo, C., Starren, E. S., Willis-Owen, S. A. G., Morris-Rosendahl, D. J., Nicholson, A. G. et al. Metabolomic, transcriptomic and genetic integrative analysis reveals important roles of adenosine diphosphate in haemostasis and platelet activation in non-small-cell lung cancer. Mol Oncol 13, 2406–2421 (2019).

16 Ruiying, C., Zeyun, L., Yongliang, Y., Zijia, Z., Ji, Z., Xin, T. et al. A comprehensive analysis of metabolomics and transcriptomics in non-small cell lung cancer. PLoS One 15, e0232272 (2020).

17 Karlsson, A., Brunnstrom, H., Micke, P., Veerla, S., Mattsson, J., La Fleur, L. et al. Gene Expression Profiling of Large Cell Lung Cancer Links Transcriptional Phenotypes to the New Histological WHO 2015 Classification. J Thorac Oncol 12, 1257–1267 (2017).

18 WHO classification of Tumours of Lung, Pleura, Thymus and Heart. 4th edn, (IARC: Lyon, 2015).

19 Lehtio, J., Arslan, T., Siavelis, I., Pan, Y., Socciarelli, F., Berkovska, O. et al. Proteogenomics of non-small cell lung cancer reveals molecular subtypes associated with specific therapeutic targets and immune evasion mechanisms. Nat Cancer 2, 1224–1242 (2021).

20 Saal, L. H., Vallon-Christersson, J., Hakkinen, J., Hegardt, C., Grabau, D., Winter, C. et al. The Sweden Cancerome Analysis Network - Breast (SCAN-B) Initiative: a large-scale multicenter infrastructure towards implementation of breast cancer genomic analyses in the clinical routine. Genome Med 7, 20 (2015).

21 Djureinovic, D., Hallstrom, B. M., Horie, M., Mattsson, J. S. M., La Fleur, L., Fagerberg, L., et al. Profiling cancer testis antigens in non-small-cell lung cancer. JCI Insight 1, e86837 (2016).

22 Li, H., Ning, S., Ghandi, M., Kryukov, G. V., Gopal, S., Deik, A. et al. The landscape of cancer cell line metabolism. Nat Med 25, 850–860 (2019).

23 Brunnstrom, H., Johansson, L., Jirstrom, K., Jonsson, M., Jonsson, P. & Planck, M. Immunohistochemistry in the differential diagnostics of primary lung cancer: an investigation within the Southern Swedish Lung Cancer Study. Am J Clin Pathol 140, 37–46 (2013).

24 Salomonsson, A., Micke, P., Mattsson, J. S. M., La Fleur, L., Isaksson, J., Jonsson, M. et al. Comprehensive analysis of RNA binding motif protein 3 (RBM3) in non-small cell lung cancer. Cancer Med 9, 5609–5619 (2020).

25 Vidarsdottir, H., Tran, L., Nodin, B., Jirstrom, K., Planck, M., Jonsson, P. et al. Immunohistochemical profiles in primary lung cancers and epithelial pulmonary metastases. Hum Pathol 84, 221–230 (2019).

26 Gyorffy, B., Surowiak, P., Budczies, J. & Lanczky, A. Online survival analysis software to assess the prognostic value of biomarkers using transcriptomic data in non-small-cell lung cancer. PLoS One 8, e82241 (2013).

27 Chong, J., Soufan, O., Li, C., Caraus, I., Li, S., Bourque, G. et al. MetaboAnalyst 4.0: towards more transparent and integrative metabolomics analysis. Nucleic Acids Res 46, W486–W494 (2018).

28 Karadzovska-Kotevska, M., Brunnstrom, H., Kosieradzki, J., Ek, L., Estberg, C., Staaf, J. et al. Feasibility of EBUS-TBNA for histopathological and molecular diagnostics of NSCLC-A retrospective single-center experience. PLoS One 17, e0263342 (2022).

29 Zang, X., Zhang, J., Jiao, P., Xue, X. & Lv, Z. Non-Small Cell Lung Cancer Detection and Subtyping by UPLC-HRMS-Based Tissue Metabolomics. J Proteome Res 21, 2011–2022 (2022).

30 Ciborowski, M., Kisluk, J., Pietrowska, K., Samczuk, P., Parfieniuk, E., Kowalczyk, T. et al. Development of LC-QTOF-MS method for human lung tissue fingerprinting. A preliminary application to nonsmall cell lung cancer. Electrophoresis 38, 2304–2312 (2017).

31 Fan, Y., Zhou, Y., Lou, M., Gao, Z., Li, X. & Yuan, K. SLC6A8 is a Potential Biomarker for Poor Prognosis in Lung Adenocarcinoma. Front Genet 13, 845373 (2022).

32 Feng, Y., Guo, X. & Tang, H. SLC6A8 is involved in the progression of non-small cell lung cancer through the Notch signaling pathway. Ann Transl Med 9, 264 (2021).

33 Loo, J. M., Scherl, A., Nguyen, A., Man, F. Y., Weinberg, E., Zeng, Z. et al. Extracellular metabolic energetics can promote cancer progression. Cell 160, 393–406 (2015).

34 Kurth, I., Yamaguchi, N., Andreu-Agullo, C., Tian, H. S., Sridhar, S., Takeda, S. et al. Therapeutic targeting of SLC6A8 creatine transporter suppresses colon cancer progression and modulates human creatine levels. Sci Adv 7, eabi7511 (2021).

35 Cancer Genome Atlas Research, N. Comprehensive molecular profiling of lung adenocarcinoma. Nature 511, 543–550 (2014).

36 George, J., Walter, V., Peifer, M., Alexandrov, L. B., Seidel, D., Leenders, F. et al. Integrative genomic profiling of large-cell neuroendocrine carcinomas reveals distinct subtypes of high-grade neuroendocrine lung tumors. Nat Commun 9, 1048 (2018).

37 Karlsson, A., Jonsson, M., Lauss, M., Brunnstrom, H., Jonsson, P., Borg, A. et al. Genome-wide DNA methylation analysis of lung carcinoma reveals one neuroendocrine and four adenocarcinoma epitypes associated with patient outcome. Clin Cancer Res 20, 6127–6140 (2014).

