## Supplementary Figure S1 for "Multi-omic profiling of squamous cell lung cancer identifies metabolites and related genes associated with squamous cell carcinoma"

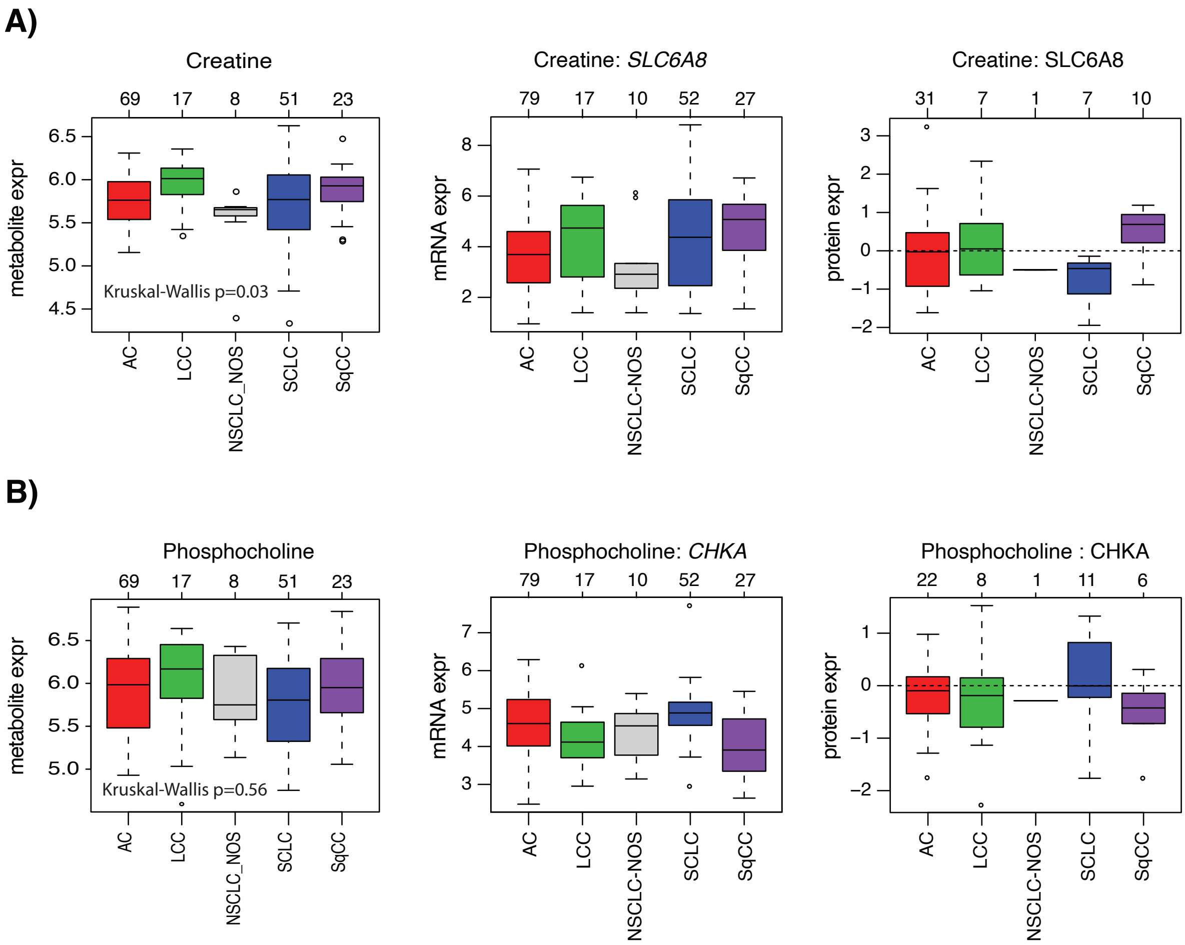


**Supplementary Figure S1.** Expression of matched metabolite, gene and protein expression of **(A)** creatine and **(B)** phosphocholine in lung cancer cell lines.
